# High-fat diet promotes Acute Promyelocytic Leukemia through PPARδ-enhanced self-renewal of preleukemic progenitors

**DOI:** 10.1101/2022.03.14.483944

**Authors:** Luca Mazzarella, Paolo Falvo, Marta Adinolfi, Giulia Tini, Elena Gatti, Rossana Piccioni, Emanuele Bonetti, Elena Gavilan Dorronzoro, Margherita Bodini, Barbara Gallo, Stefania Orecchioni, Bruno Achutti Duso, Chiara Ronchini, Anna Giulia Sanarico, Rani Pallavi, Sophie Roerink, Francesco Bertolini, Myriam Alcalay, Gaetano Ivan Dellino, Pier Giuseppe Pelicci

**Affiliations:** IRCCS European Institute of Oncology, Milan, Italy; University of Seville, Department of Biochemistry and Molecular Biology, Seville, Spain; Wellcome Trust Sanger Institute, Hinxton Cambridgeshire UK; Department of Hemato-Oncology, Universita’ Statale di Milano

**Author notes:** Correspondence to: Luca Mazzarella MD, PhD, and Pier Giuseppe Pelicci MD PhD, Address: Department of Experimental Oncology, Via Adamello 16, 20141 Milan, Italy. These authors have contributed equally (Luca Mazzarella, Paolo Falvo).

## Abstract

Obesity is associated with a higher risk of developing many cancer types including acute promyelocytic leukaemia (APL), a subset of acute myeloid leukemias (AML) characterized by expression of the PML-RARα oncogene. The molecular mechanisms linking obesity and APL development are not known. To model clinical observations, we established a mouse model of diet-induced obesity using transgenic mice constitutively expressing PML-RARA α in the hematopoietic system (PML-RARα KI mice) fed either standard (SD) or high-fat (HFD) diets. HFD-fed PML-RARα KI mice developed leukaemia with reduced latency and increased penetrance, as compared to SD-fed mice. HFD leads to accumulation of DNA damage in hematopoietic stem cells (HSCs), but, surprisingly, this was not associated with mutational load gain, as shown by whole genome/exome sequencing of pre-leukemic and leukemic cells. Importantly, very few of the observed mutations were predicted to act as cancer drivers, suggesting the relevance of nongenetic mechanisms. HFD led to an expansion of hematopoietic progenitor cells with a concomitant reduction in long-term hematopoietic stem cells, and in the presence of PML-RARα this was also accompanied by an enhancement of *in vitro* and *in vivo* self-renewal. Interestingly, Linoleic Acid (LA), abundant in HFD, recapitulates the effect of HFD on the self-renewal of PML-RARα HPCs by activating the peroxisome proliferator-activated receptor delta (PPARδ), a central regulator of fatty acid metabolism involved in the promotion of cancer progression. Our findings have implications for dietary or pharmacological interventions aimed at counteracting the cancer-promoting effect of obesity.

**Key points:**

- high fat diet (HFD) promotes APL leukemogenesis in mouse models, reproducing the exquisite sensitivity to obesity observed in humans
- although HFD leads to DNA damage and mutations, the molecular mechanism is nongenetic and linked to the transcription factor PPARδ

## Introduction

Despite overwhelming evidence and the growing size of the obesity epidemic, our understanding of the molecular mechanisms underlying the correlation between obesity and cancer, including hematological neoplasms^1,2^ is remarkably poor, partly because of the difficulty in modelling the pleiotropic effects induced by diet with appropriate reductionist approaches. For instance, it is unknown whether obesity intervenes in the early or late phases of cancerogenesis, whether it impacts on genetic or nongenetic mechanisms of cancer promotion, and the relative contribution of specific dietary components or endogenous hormones and inflammatory cytokines produced by the increased fat tissue ^1^. Although it is likely that a combination of multiple factors intervenes at different stages, identifying the precise role of each contributing factor may provide the rationale for informing more specific policies aimed at preventing and treating obesity-associated cancers. We recently reported that among hematological malignancies, risk and outcome of Acute Promyelocytic Leukemia (APL) are strongly affected by Body Mass Index (BMI)^3,4^.

The t(15;17) translocation, leading to the PML-RARα fusion, is considered the initiating genetic event for most APLs, but current theory posits that additional genetic or nongenetic events are required to generate a full blown leukemia ^5–8^. APL mouse models faithfully recapitulate clinical-biological features of the disease, including the typical promyelocytic morphology of leukemic blasts and its sensitivity to APL-treating agents (e.g.; all-trans retinoic acid (ATRA) and arsenic trioxide) ^9–11^. In the widely used mouse APL model developed by Ley et al, the human PML-RARα is knocked-in under the control of the mouse cathepsin G gene (PML-RARα knock-in, PRKI) to allow specific expression of the fusion protein in the mouse hematopoietic compartment ^11^. This model is particularly appropriate for studying the events preceding leukemia (pre-leukemic phase), as mice develop full-blown leukemia with a long latency and incomplete penetrance, thus allowing analyses of the effects on leukemogenesis elicited by environmental cues such as high-fat diet (HFD), an established model of diet-induced obesity and metabolic stress ^12^. Taking advantage of this model, we investigated molecular mechanisms underlying the effect of HFD on APL leukemogenesis.

## Results

### HFD enhances leukemogenesis in PML-RAR knock-in mice

PRKI mice were randomized at 8-10 weeks of age to receive normal chow (Standard Diet; SD) or a diet with 60% animal fats (High Fat Diet; HFD). Weight gain was equal in wild type (WT) and PRKI mice (figure 1A). In line with prior studies, PRKI mice fed with SD had incomplete disease penetrance at ∼70% and a median Leukemia Free-Survival (LFS) of 254 days. HFD increased dramatically disease penetrance to 100% and accelerated median LFS to 204 days (p<0.001). There was no effect of HFD alone on the mortality of WT mice (Figure 1B). Necroscopy of moribund mice revealed that SD and HFD mice developed a disease similar to that in SD mice, with splenomegaly, loss of splenic architecture, liver infiltration and blast cells in the peripheral blood (figure 1D-E). Thus, diet-induced obesity accelerates APL onset in this mouse model, suggesting the existence of specific and conserved molecular mechanisms through which obesity promotes APL leukemogenesis.

**Figure 1.**
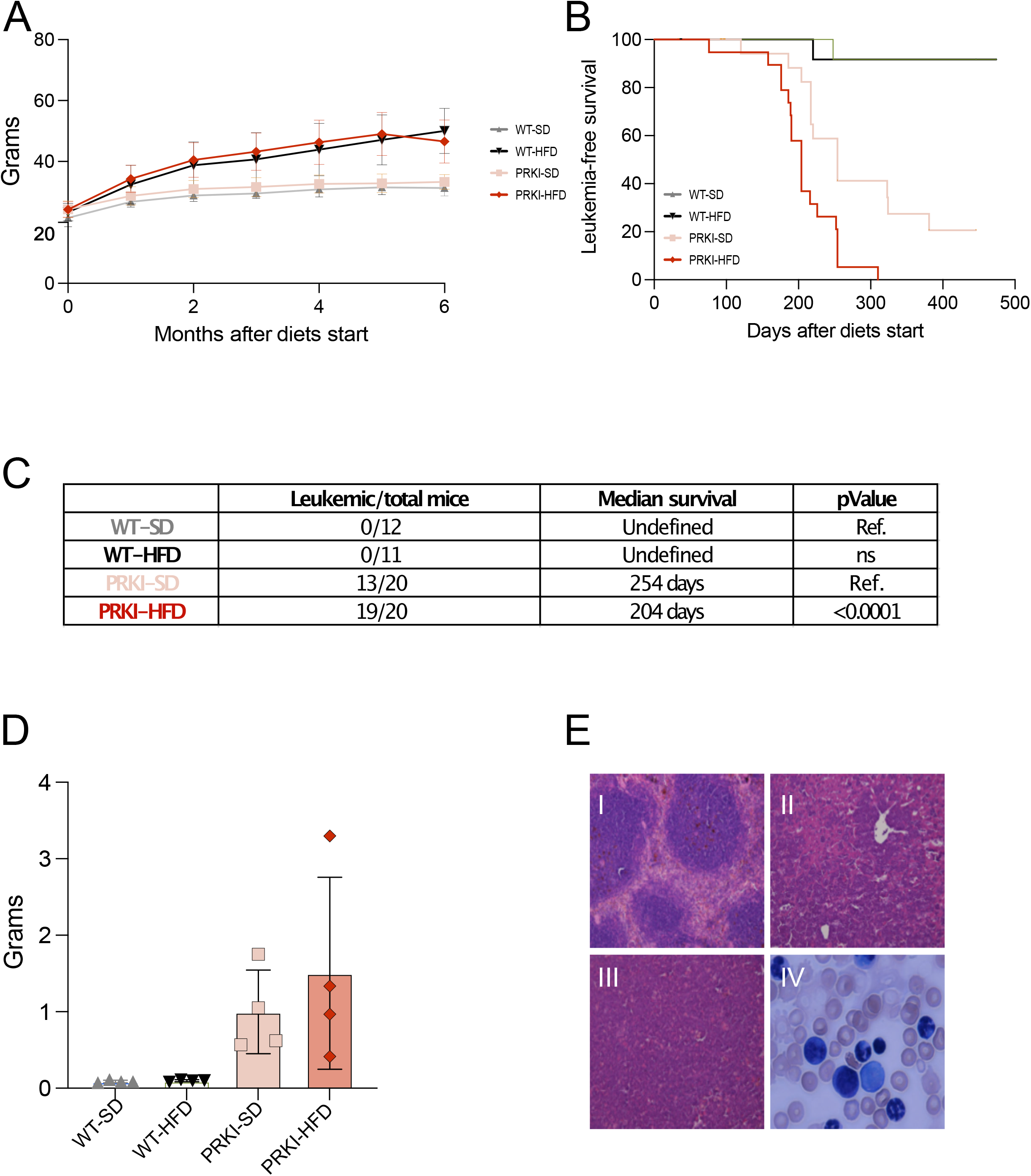
High Fat Diet promotes PML-RARA leukemia development. **A**. Weight measurement of WT and PRKI mice after the initiation of SD or HFD. Weight gain was identical in both strains. **B**. Kaplan-Meier curve of leukemia-free survival (LFS) after diet start in WT and PRKI mice fed HFD or SD as indicated. NB 1 mouse per each group in the PRKI/HFD, WT/SD and WT/HFD groups died due to non-leukemic reasons (cannibalism) and were censored; 1 mouse per diet group in WT mice died at approximately one year of age in poorly defined circumstances and was not censored. **C** Table of relevant statistics relative to exp 1A-B. P value as per log-rank test against genotype-matched SD (“ref”) **D-E** results of necroscopy of four mice per group obtained upon sacrifice for humane reasons (PRKI) or at experiment termination (WT). **D**. spleen weight. **E** representative images of hemopoietic tissues: **I**) spleen of an aged WT/SD animal, with clearly distinguishable red/white pulp (100x); **II**) spleen of a HFD/PRKI mouse, with complete architecture subversion and infiltration by leukemic blasts (100x); **III**) liver from the same mouse, with extensive periportal infiltration; **IV**) peripheral blood smear from the same mouse, showing blasts and several abnormally segmented granulocytes (100x)

### HFD causes oxidative DNA-damage in pre-leukemic stem cells

Accumulation of oxidative DNA damage has been reported in lymphocytes of obese people ^13^ and suggested to be connected with the development of obesity-related diseases, including cancer ^1,14^. DNA damage can lead to an increased rate of DNA mutations, thus affecting tumor onset and progression ^13^. Thus, we first tested if HFD-accelerated leukemogenesis is associated with accumulation of oxidative DNA damage. We measured DNA strand breaks (single and double) using the alkaline comet assay on FACS-isolated HSCs (defined as Lin-Sca+ c-Kit+ Cd150+ Cd48-). In both WT and PRKI mice, HFD increased significantly the percentage of DNA damage, to an extent that was slightly lower than that acutely induced by UV irradiation (Figure 2A-B). Comet analyses of more committed subpopulations (progenitors sorted as Lin-Sca+ c-Kit+ and depleted of Cd150+Cd48- or differentiated cells sorted as lineage-positive; Lin+) in PRKI mice revealed that accumulation of HFD-induced DNA damage is progressively less-pronounced in more differentiated cells (Figure 2C), confirming that DNA damage is repaired along with cell differentiation, as previously reported ^15,16^. Remarkably, administration of the antioxidant n-Acetylcysteine (NAC) in the drinking water completely abolished HFD-induced DNA damage in comet assays of PRKI HSCs, demonstrating the oxidative origin of HFD-induced DNA damage in these cells (Figure 2D). We conclude that prolonged HFD induces accumulation of oxidative DNA-damage in HSCs.

**Figure 2.**
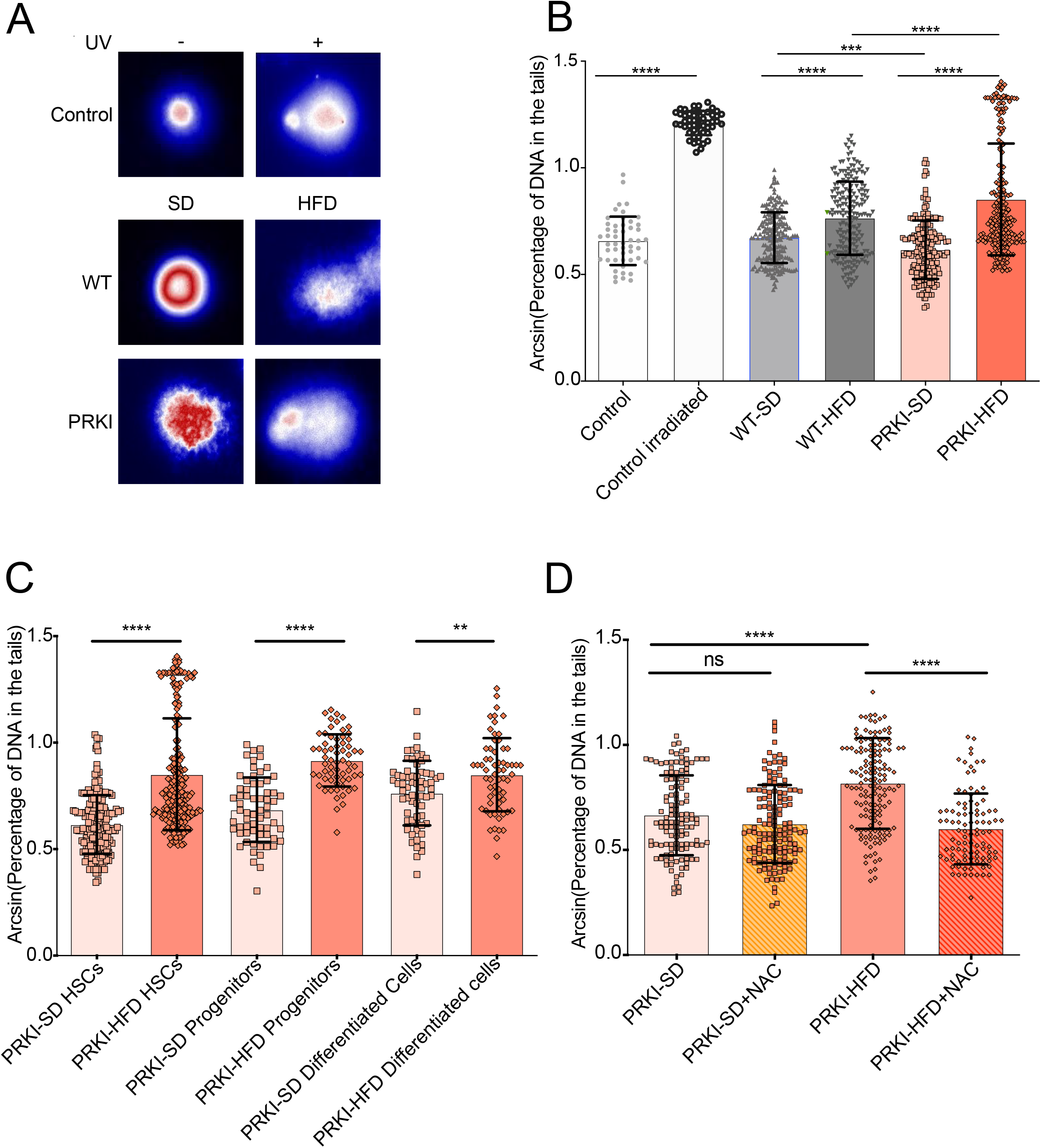
High Fat Diet increases oxidative DNA damage in HSCs. **A-B**. Representative images (A) and quantification (B) of comet assay on HSCs in irradiated/unirradiated controls and WT SD, WT HFD, PML-RARα KI SD, and PML-RARα KI HFD-fed mice. Percentage of DNA in the comet tails are shown after arcsine-transformation to better appreciate differences at the extreme ends of the distribution. Due to non-normal distribution in some samples (assessed by Shapiro-Wilk’s test), a nonparametric 2-tailed Mann-Whitney test was used to assess pairwise differences (*p<0.05; ***p<0.001; ****p<0.0001). n=5 mice per condition, for each condition at least 40 events were acquired; for controls 1 mouse per condition was used and 50 events acquired. **C)** Quantification of comet assay on HSCs, progenitors and differentiated cells expressed as arcsin of percentage of DNA in the tails in PML-RARAα KI SD and PML-RARA α KI HFD fed mice. Data were analysed using Shapiro-Wilk’s test to assess normal distribution, 2-tailed Mann-Whitney test for non-normal distribution to assess difference between groups (***p<0.001; ****p<0.0001). n=5 mice per condition, for each condition at least 40 events were acquired. **D)** Quantification of comet assay on HSCs expressed as arcsin of percentage of DNA in the tails in PML-RARα KI SD and PML-RARα KI HFD, in PML-RARα KI SD+NAC, and PML-RARα KI HFD+NAC treated mice. Data were analysed using Shapiro-Wilk’s test to assess normal distribution, 2-tailed Mann-Whitney test for non-normal distribution to assess difference between groups (****p<0,0001). n=4 mice per groups, and at least 40 events acquired per mouse.

### HFD is not associated with increased leukemia driver-gene mutations in pre-leukemic or leukemic cells

Accumulation of DNA damage in HSCs suggests that HFD may promote leukemogenesis by enhancing the likelihood of additional key genetic events in pre-leukemic cells. Analysis of the TCGA cohort of 16 APLs ^3,8^ showed no strong correlation between BMI and age-normalized mutations (Figure 3A-B). However, this analysis was poorly informative due to small sample size and imbalanced BMI classes (only 2 patients in the TCGA have a normal BMI).

**Figure 3.**
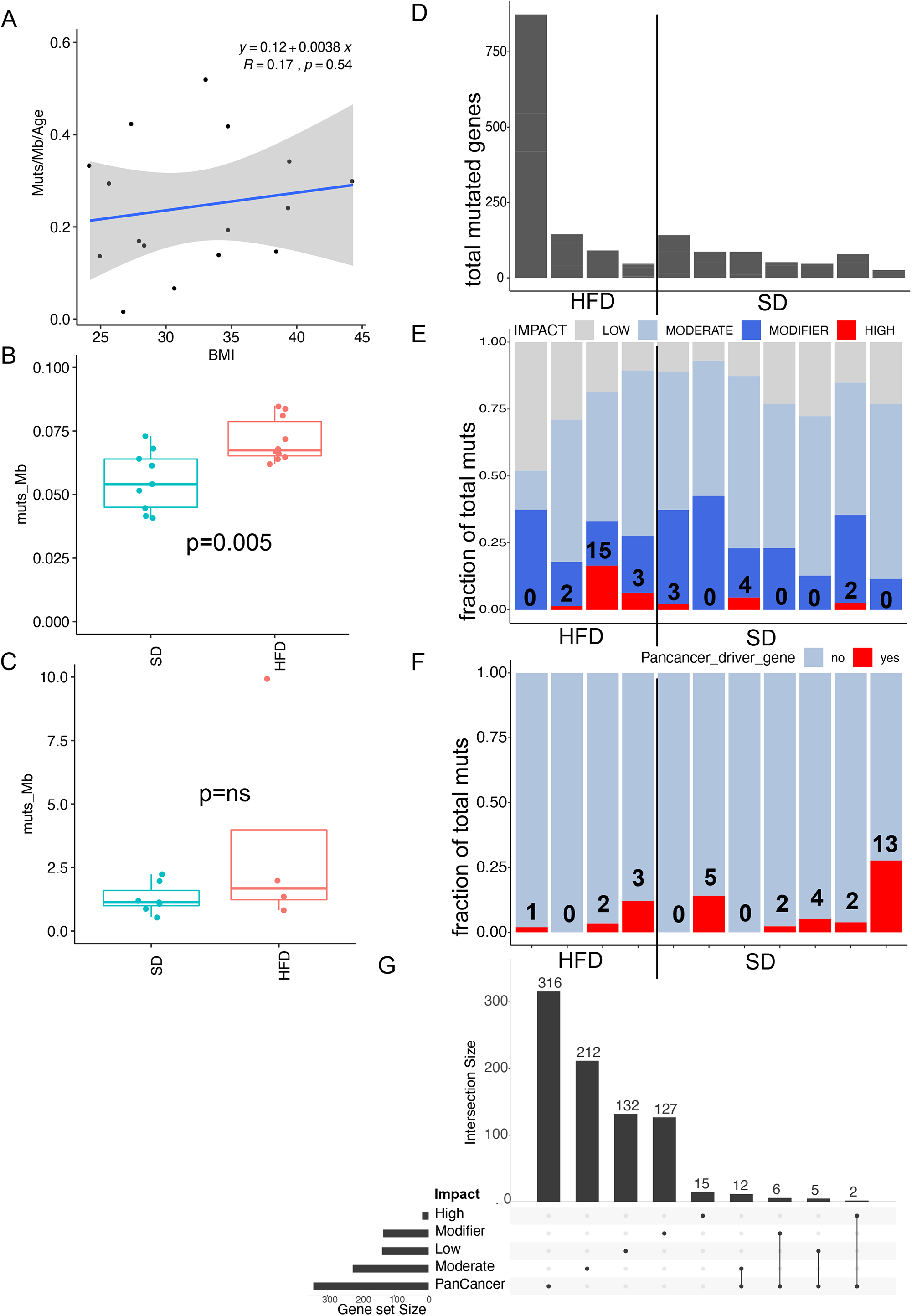
Obesity and HFD is associated with higher mutation frequency in APL and HSC but has minimal impact on driver gene mutations. **A** correlation between BMI and age-normalized mutations in APLs from the TCGA dataset. The bar shows the linear regression (equation shown on chart), grey area shows 95% confidence interval. **B, C**. Mutations per Megabase in mouse HSC colonies (in B) and APLs (in C) by diet. P values are calculated by 2-tailed t-test **D-F**. Analysis of functional consequences of mutations in APL samples. Each column represents a single sample from figure C D. Total number of mutated genes. E. VEP analysis of functional impact of mutations; superimposed numbers indicate number of high-impact mutations; F. Mutations involving (superimposed numbers) or not a PanCancer Driver gene G. Intersection between different VEP and PanCancer classes in the entire dataset.

To directly measure the impact of diet on genetic events in our experimental system, we sequenced the genome of leukemic and pre-leukemic cells in PRKI mice subjected to SD or HFD. In established leukemias, in which driver and passenger mutations are present at relatively high allele frequency, we analysed bulk populations by Whole Exome Sequencing (WES), searching for specific genes or pathways preferentially mutated in HFD. For preleukemic cells, which have not yet undergone clonal selection and expansion, we sought to quantitate genomic mutations by single-cell whole-genome sequence, using clonal populations grown from single HSCs sorted from HFD or SD-fed PRKI pre-leukemic mice. Tail WGS was used as internal reference to call mouse-specific somatic variants.

In both established APLs and pre-leukemic HSC-derived colonies, HFD was associated with a moderate increase in the number of mutations per megabase (mut/Mb), which was statistically significant in the preleukemic colonies but not in APLs, where variance was wider (figure 3D). Larger structural abnormalities like copy number variants and translocations were not significantly different (supplementary figure 1 and supplementary table 1). Loss of the histone demethylase Kdm6a, previously implicated in the acquisition of the leukemic phenotype in the PMLRAR mouse model ^17^, was confirmed in the majority of samples in this dataset (supplementary table 2), without differences between diets; no recurrent alteration could be identified To understand if this moderate increase in somatically acquired mutations actively contributes to HFD-enhanced leukemogenesis, we analysed their functional impact by the Ensembl Variant Effect Predictor (VEP) ^18^ and assessed whether the affected genes are among established cancer drivers identified by the PanCancer project ^19^. No statistically significant difference between groups was identified, with HFD bearing nominally more mutated genes in total (mean 289 vs 74, figure 3D) and more VEP “high impact” genes (mean = 5 vs 1.3, figure 3E), but fewer mutated PanCancer driver genes (mean=1.5 vs 3.7, figure 3F) than SD. Considering all mutated genes in the dataset together, only 2 genes were both “high impact” by VEP and included among PanCancer drivers: XPO1 and PIK3CB (figure 3G). However, both were both affected by loss-of-function nonsense mutations (supplementary table 3), which makes them unlikely to drive leukemogenesis in our system since they have been reported as gain-of-function oncogenes ^20,21^.

Collectively, these results confirm that the frequency of genetic events in both APL preleukemic and leukemic cells is very low and is, at most, only slightly increased upon exposure to HFD. The additional mutations observed in HFD do not appear to impact *bona fide* leukemia driver genes, suggesting that genetic events are unlikely to play a causative role during HFD-induced leukemogenesis, prompting us to investigate non-genetic mechanisms.

### HFD expands the pool of preleukemic progenitors without loss of self-renewal in the presence pf PML-RARA a

The differentiation block imposed by PML-RARα is considered a critical component of the leukemia phenotype in APLs, since reversion of PML-RARα activity by high-dose ATRA inhibits self-renewal, releases the differentiation block and leads to disease remission in most cases ^22,23^. Thus, we investigated if HFD regulates self-renewal and differentiation of myeloid progenitors in a PML-RARα-dependent manner.

We first investigated the distribution of stem and progenitor cells in the BM of WT and PRKI mice fed SD or HFD for 4 months. The LSK compartment was divided into HSCs (LSK d150+CD48-), multipotent hematopoietic progenitors (MPP: LSK CD150-CD48-) and restricted progenitors HPC-1 (LSK CD150-CD48+) and HPC-2 (LSK CD150+ CD48+)25. HFD had a similar impact on both PRKI and WT mice, significantly increasing both relative and absolute numbers of committed HPC1 progenitors, at the expenses of HSCs (Figure 4A-B), in agreement with previous studies on WT mice ^24^. Notably, abrogation of oxidative DNA damage by simultaneous administration of NAC did not abrogate HFD-induced HSC loss and HPC expansion; perhaps paradoxically, NAC seemed to perturb hematopoiesis in the same way as HFD. Though the mechanism of this phenomenon was not investigated further, these data clearly imply that oxidative DNA damage (figure 2x) is not per se the trigger of HFD-induced HPC expansion in PML-RARα expressing cells.

**Figure 4.**
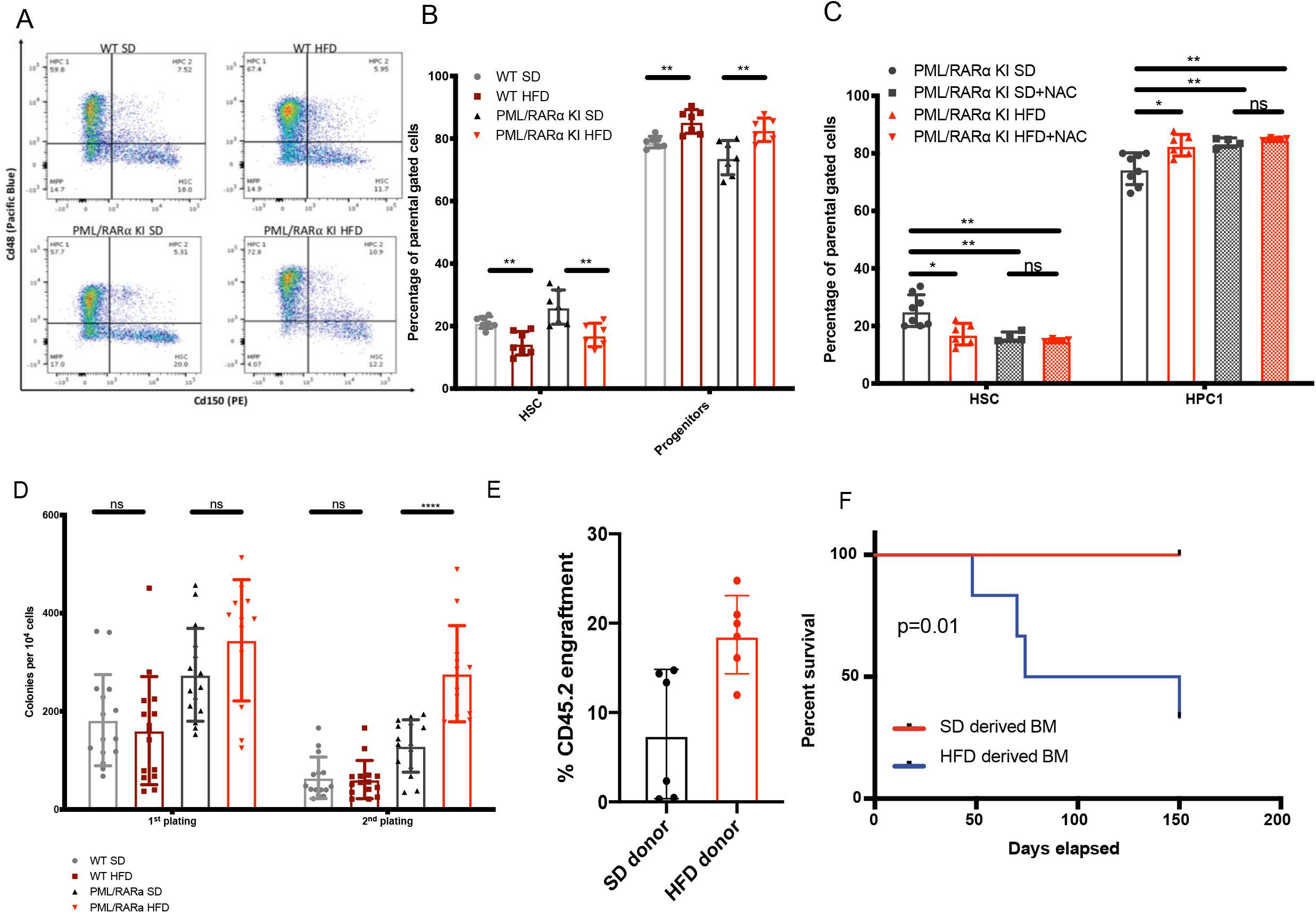
High fat Diet increases self-renewal in PML-RARA α KI mice. **A**. Representative FACS plot of HSCs and progenitors derived from WT and PRKI mouse fed with SD or HFD. Gating strategy used is extensively explained in25. Briefly, cells were gated for morphological parameters to discriminated alive cells and to exclude cell doublets. Cells were than analysed for Lin positivity, and Lin- cells were gated. Gated Lin- cells were analysed for Sca+ and c-Kit+ positivity (LSK cells). LSK cells were finally analysed for CD150 and CD48 positivity. The plot shows positivity of LSK cells for CD150 and CD48 and four subpopulations were discriminated: HSCs (Cd150+Cd48-), MPP (Cd150-Cd48-), HPC1 (Cd150+Cd48-), HPC2 (Cd150+Cd48+). Percentage of cells in each gated subpopulation is indicated in the plot. **B**. Quantification of FACS analysis shown in A. Percentage of cells divided for genotypes and diets in the four stem cell populations: HSCs (Cd150+Cd48-), MPP (Cd150-Cd48-), HPC1 (Cd150+Cd48-), HPC2 (Cd150+Cd48). Data were analysed using Shapiro-Wilk’s test to assess normal distribution, 2 tails unpaired t test for normal distribution to assess difference between groups (*p<0.05; **p<0.01). n=8 per each experimental group. **C**. Effect of NAC on stem/progenitor cells. Data were analysed using Shapiro-Wilk’s test to assess normal distribution, 2 tails unpaired t test for normal distribution to assess difference between groups (*p<0.05; **p<0.01). n=8 for SD and HFD groups; n=4 for NAC treated groups. **D**. Colonies count per 10^4^ total BM cells plated in M3434 at two different passages. BM cells from WT and PML-RAR KI mice fed with SD or HFD were collected and 10^4^ cells were plated in M3434. After one week, colonies were counted (first plating) and disaggregated to obtain single cell suspension. Next, 10^4^ single cells were plated in M3434 and counted again after one week (second plating). Two-tailed unpaired t-test was performed (**p<0.01; ****p<0.0001). n=4 mice per experimental group and for each mouse BM cells were plated in triplicate. **E-F**. One-month engraftment of Cd45.2 cells derived from total BM of SD- and HFD-fed PML-RARα KI mice (2 pooled donors per diet) in twelve Cd45.1 WT recipient mice (6 per donor pool). Data were analysed using Shapiro-Wilk’s test to assess normal distribution, 2 tails unpaired t test for normal distribution to assess difference between groups (**p<0.01). **G**. Kaplan-Meier curve of Cd45.1 mice transplanted with total BM derived from Cd45.2 PML-RARA α mice fed with SD or HFD. Each vertical step in the curve indicates one or more events (i.e.; deaths). On y axis is reported the percentage of leukemia free survival while on x axis time after experiment starts. Mantel-Cox test was performed (**p<0.01). n=6 per experimental group.

The HFD-expanded HPC compartment is associated with decreased self-renewal in wt mice ^24,25^. However, this is in contradiction with the observed leukemia-enhancing effect of HFD, since the founding leukemia cells is expected to possess unlimited self-renewal ability. Thus, we directly tested diet-induced changes in self-renewal by performing *in vitro* colony-forming assays (to test short-term self-renewal of committed progenitors) and *in vivo* competitive transplantation assays (to test long-term self-renewal of stem and progenitor cells), using total BM from SD or HFD-fed WT or PRKI mice. In the colony formation assay, efficiency upon secondary replating was reduced in WT mice and was higher in PRKI mice, as previously reported ^23^. However, HFD had no effect on primary and secondary plating efficiency in WT BM cells, whereas it significantly enhanced secondary replating of PRKI BM cells. For competitive transplantation assay, BM cells from SD or HFD-fed PRKI mice (expressing the Cd45.2 allelic variant) were mixed at 1:9 ratio with wild type BM cells (expressing Cd45.1) and transplanted into irradiated Cd45.1 recipient mice, which were then maintained on a SD. Recipients of HFD-fed donors were more efficiently engrafted than recipients of SD donors after 1 month (median=19.2% CI 95% [14.1-23.3%] vs 7.9%, CI 95% [0-15%], p=0.015) (Figure 4D). Strikingly, all recipients of HFD PRKI cells succumbed to leukemia in the first transplant (median survival 74 days vs 150, p=0.01) (Figure 4E). Together, these data demonstrate that HFD increases significantly self-renewal and repopulating capacities of PML-RARA α BM cells, and that acquisition of the leukemogenic phenotype is cell-intrinsic and maintained even upon removal of the HFD environment upon methylcellulose serial replating in or transplant into SD mice.

### Linoleic Acid reproduces the effect of HFD on self-renewal *in vitro*

We previously reported the selective upregulation of polyunsaturated fatty acid metabolism in APLs, as compared to other AMLs ^3^. Linoleic Acid (LA) is enriched in high fat western diets and is one of the major components of the most widely used mouse HFD. Thus, we tested if mono- (oleic, OA) or poly- (e.g., LA and arachidonic acid, AA) unsaturated fatty-acids replicate the enhanced self-renewal observed in HFD-fed PRKI BM. In colony formation assays, LA significantly increased replating efficiency at later passages (Figure 5A), whereas its derivative AA had a generalized toxic effect that prevented colony growth, event at concentrations ∼10 fold lower than LA; the mono-unsaturated oleic acid had a modest effect compared to LA. LA administration was required continuously for its effect on self-renewal, since its removal during passaging reverted its effect, leading to significantly less-efficient colony formation (Figure 5B). Finally, we assessed the effect of LA on self-renewal of selected stem/progenitors populations, isolated ex vivo through FACS sorting. Serial replating of HSC, MPP, HPC1 and HPC2 from PRKI BM showed that enhanced LA-induced self-renewal on HPCs, but not on HSCs nor MPPs (Figure 5C). Thus, in the presence of PML-RARA a, HPCs are the sub- population most expanded by HFD and the most sensitive to the self-renewal-promoting effect of LA.

**Figure 5.**
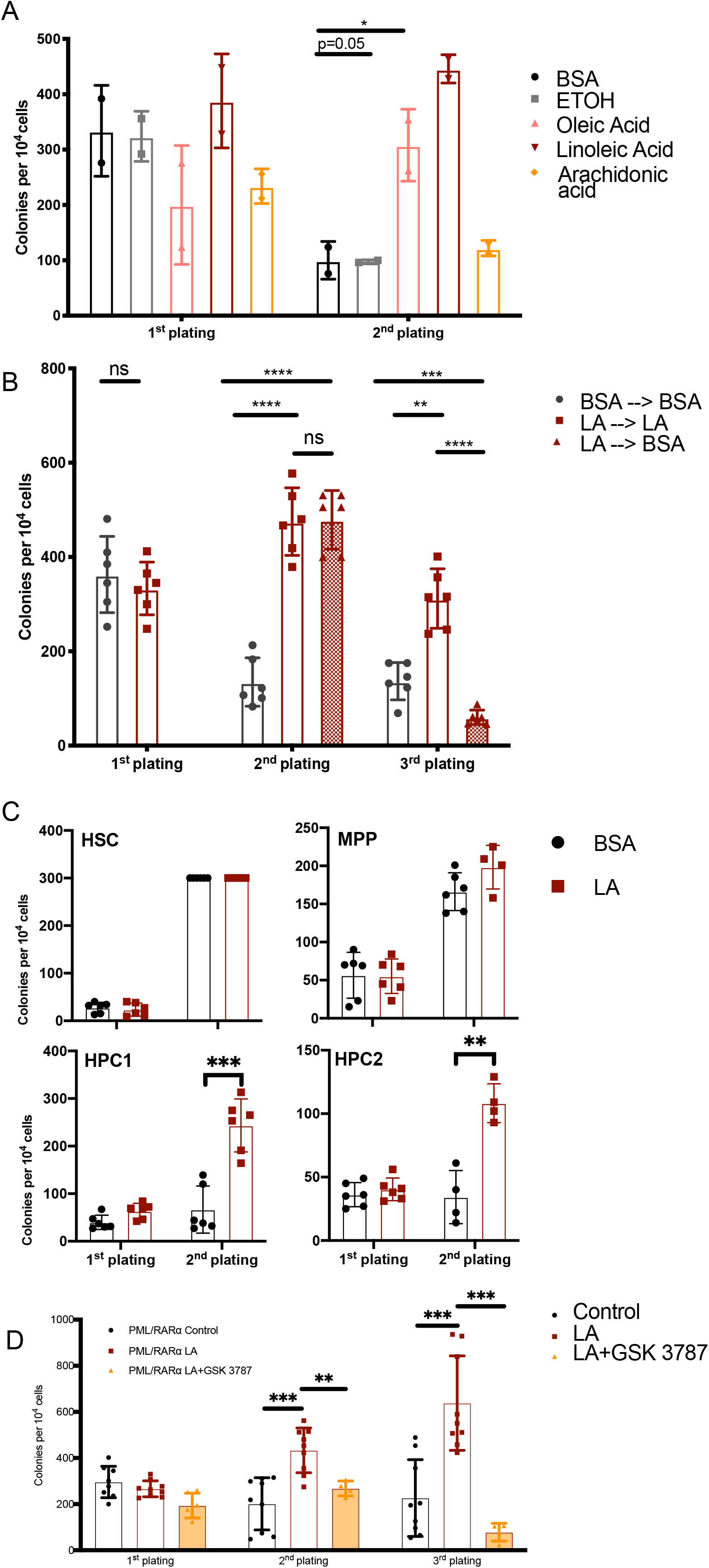
LA enhances self-renewal activity in PML-RARα KI HSCs. **A**. Colonies count per 10^4^ total BM cells from PRKI mice serially replated in M3434, in presence of BSA+EtOH as control, or OA, LA, AA. Two-tailed unpaired t-test (**p<0.01). n=2 mice per experimental group and for each mouse BM cells were plated in triplicate. **B**. Colonies count per 10^4^ total BM cells from PRKI mice serially replated in M3434, in presence of BSA+EtOH as control, or OA, LA, AA. Two-tailed unpaired t-test (***p<0.001; ****p<0.0001). n=2 mice per experimental group and for each mouse BM cells were plated in triplicate. **C**. Colonies count upon plating of hematopoietic stem cells (HSC), or hematopoietic progenitor cells 1 (HPC1), hematopoietic progenitor cells 2 (HPC2) and multipotent progenitors (MPP) in M3434 in presence BSA or LA. Two-tailed unpaired t-test (**p<0.01; ***p<0.001). n=3 mice per experimental group and for each mouse BM cells were plated in duplicate. Counts are expressed per 10^4^ cells for all cell types except HSC, which are expressed per 100 cells (1^st^ plating) and per 1000 cells (second plating), due to massively different yield in the two platings BSA, bovine serum albumin; EtOH, ethanol; OA, oleic acid; LA, Linoleic Acid; AA, arachidonic acid.

### LA activity depends on PPARδ activation

To gain insight into the biological function of LA on APL cells, we analysed transcriptional changes induced by LA treatment of the NB4 APL-model cell line, by RNAseq. LA resulted in the differential regulation of 168 genes (25 upregulated and 143 downregulated, figure 6A and supplementary table 4). Functional analyses through GSEA revealed a clear modulation of several lipid metabolism pathways, with up-regulation of oxidative-phosphorylation pathways and downregulation of steroid biosynthesis (figure 6B) and other lipid biosynthetic pathways (supplementary table 5 for complete list). In particular, genes encoding for the two key enzymatic steps of FAO, the Carnitine palmytoil transferase 1 (CPT1A) and the fatty acid uptake carrier SLC25A20, were upregulated in NB4 cells by LA (figure 6C), consistent with a metabolic switch towards fatty-acid oxidation, a bioenergetic pathway previously shown to be associated with increased survival and resistance to apoptotic stimuli in leukemic cells ^26^. Interestingly, KEGG enrichment analysis of differentially-regulated genes also showed, among others, a significant enrichment for genes involved in peroxisome proliferator-activated receptors (PPAR) signaling (figure 6D and supplementary table 6), suggesting that LA functions by directly modulating PPAR transcriptional-activity. Notably, LA is transformed by intracellular lipoxygenases into endogenous activators of PPARs, in particular of PPARα and PPARδ ^27,28^.

**Figure 6.**
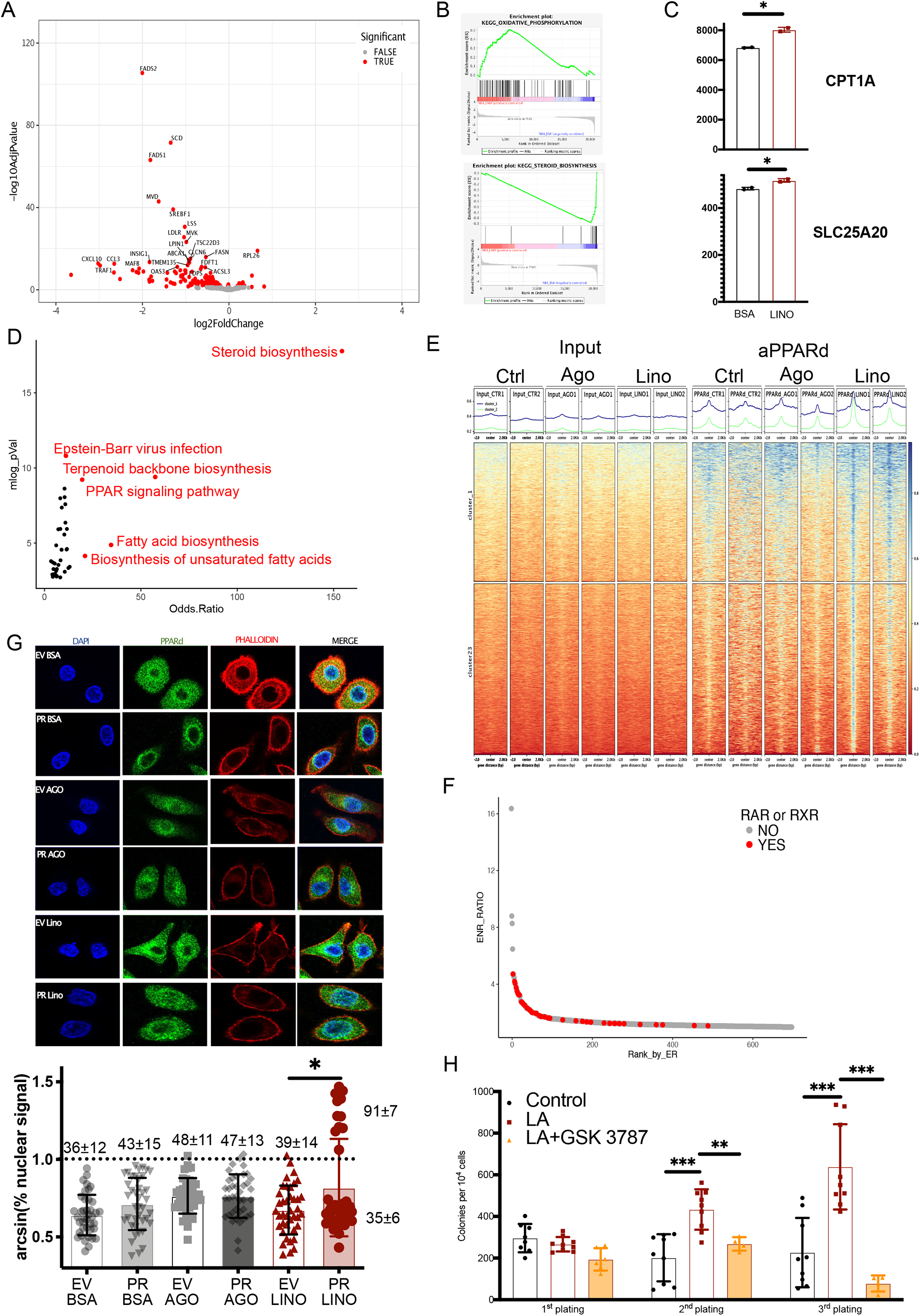
Linoleic treatment triggers PPARδ translocation in the nucleus and activation of specific transcriptional program. **A**. Volcano plot showing differential gene expression between 5 mM LA- and BSA- (vehicle) treated NB4. Red dots are genes which are statistically significant (adjusted pvalue <0.01). **B**. KEGG enrichment (Odds ratio vs -log10 pVal) of differentially expressed genes between LA- and BSA- (vehicle) treated NB4 cells. Highlighted in red are gene sets with Odds Ratio >= 10 and adjusted pVal <10^−7^ **C**. Expression (RPKM) of key genes in FAO **D**. Expression (RPKM) of PPAR family members **E**. heatmap and unsupervised clustering of 3796 PPARδ ChIPseq peaks in NB4 cells treated with BSA, PPARδ agonist (GW501516) or LA, comparing input vs IP signal **F**. Motif enrichment analysis. Significantly enriched motifs are plotted in descending order by Enrichment ratio (ER) highlighting high prevalence of known RAR or RXR-binding motifs among the top enriched PPARδ-bound motifs **G-H**. Quantification of PPARδ translocation in the nucleus by IF for PPARδ and phalloidin/DAPI to identify nucleus and cell borders for EV and PML-RARA α transduced HeLA cells treated with BSA, PPARδ agonist (GW501516) or LA; in G, representative images; in H, nuclear signal quantification after arcsine-transformation of percentage nuclear signal to better appreciate differences at the lower end of tail percentage. Overlayed numbers indicate average percentage ± standard deviation; distributions were normal by Shapiro-Wilk’s test for all sets except for PMLRAR-LINO, so statistical significance was assessed by Fisher’s test, considering the number of cells that were above vs below the maximum background level computed in ctrl-BSA (dashed line) (*p<0.05). n=60 events per experimental condition. **I**. Serial colony formation assay of PRKI BM cells exposed to BSA (vehicle) or LA o LA + GSK3787 (PPARδ inhibitor). Data were analysed using Shapiro-Wilk’s test to assess normal distribution, 2 tails unpaired t test for normal distribution to assess difference between groups 2 tails unpaired t-test was performed (***p<0.001). n=3 mice per experimental group and for each mouse BM cells were plated in triplicate for control and LA, and duplicate for LA and GSK3787.

PPARδ regulates expansion and self-renewal of normal HSCs and intestinal cancer SCs in response to HFD ^29,30^, and is expressed at high levels in NB4 cells, as compared to the other members of the PPAR family (supplementary figure 2A). Thus, we analysed genome-wide binding of PPARδ by Chromatin Immunoprecipitation-sequencing (ChIPseq) upon treatment of NB4 cells with vehicle, LA or the PPARδ agonist GW501516. We identified 3796 sites that were significantly bound in at least one of the three conditions (supplementary table 7). Both GW501516 and LA significantly increased PPARδ binding signal, as compared to controls; hierarchical clustering identified two groups of PPARδ sites or peaks: 1738 in common between GW501516 and LA treatments (cluster 1), 2058 specific for LA (cluster 2), but no GW501516-specific peak-clusters (figure 6E). PPARδ peaks were enriched in intronic and intergenic areas (supplementary figure 2B) and mostly located >10 kb from the nearest TSS, a distribution that rendered difficult the univocal association between peaks and regulated genes. Nevertheless, limiting analyses to peaks occurring within 2.5 kb from the nearest transcription start site (TSS) allowed identification of 351 *bona fide* direct PPARδ direct target-genes (supplementary table 7). Functional enrichment analysis by *enrichr* found, as expected, an enrichment for genes known to be regulated by PPAR transcription factors and involved in the regulation of lipid metabolism, in particular fatty acid beta oxidation, lipid transport and respiratory electron transport (supplementary figure 2C and supplementary table 8), coherently with transcriptomic data. Motif enrichment analyses of all the 3796 PPARδ peaks showed strong enrichment for known binding-motifs of the obliged cofactor RXR and, most importantly, of various Retinoic Acid Receptor Elements (RARE), the DNA-binding consensus sequence of RAR and PML-RARα ^31^, suggesting co-binding with PML-RARα in these cells (figure 6F and supplementary table 9). To formally test if PMLRARα favors LA-induced PPARδ-binding to chromatin, we assessed nuclear-cytoplasmic shuttling of PPARδ by immunofluorescence (IF), in response to LA or the PPARδ-agonist in HeLa cells expressing PMLRARα or empty vector (EV) (figure 6G). In EV, control-treated cells, ∼36% of the PPARδ signal was found in the cytoplasm, with no cells exhibiting >60% signal in the nucleus. While addition of the PPARδ-agonist and/or infection with PMLRAR caused a homogeneous, moderate, yet non statistically-significant increase in signal; LA induced a clearly bimodal distribution with the emergence of a clearly distinct population with prevalent nuclear signal. Collectively, these experiments suggest that LA drives cooperation between PMLRARα and PPARδ at the chromatin level to regulate the transcription of genes involved in lipid metabolism. To directly test the relevance of this circuit for leukemia promotion, we assessed the effect of the selective PPARδ inhibitor GSK3787 ^32^ on the self-renewal capacity of PMLRARα bone marrow cells in serial colony-formation assays. As shown in figure 6H, simultaneous exposure to GSK3787 and LA significantly reduced colony formation at late passages, completely abrogating LA-enhanced self-renewal. Thus, we conclude that PPARδ is a key mediator of the leukemia-promoting activity of HFD and LA in the presence of PMLRARα.

## Discussion

In this study we provide mechanistic evidence that obesity directly promotes APL leukemogenesis through a PPARδ/PMLRARα transcriptional circuit that is activated by PUFA and in particular LA. This leads to the outgrowth of activated and self-renewing HSPCs through nongenetic mechanisms.

Our data confirm a previous study showing that HFD expands the pool of committed HPCs, a process that is accompanied by stem/progenitor cell exhaustion ^24^. In the presence of PML-RARα, instead, HFD-activated HPCs do not exhaust and acquire sustained self-renew, both *in vitro* and *in vivo*, leading to leukemia in the long run. Thus, the functional interaction between HFD and PML-RARα leads to a net expansion of the pool of preleukemic myeloid progenitors ^23,33^.

We identified a specific signaling circuit in which the high-fat dietary-component Linoleic Acid (LA) activates directly the transcription factor PPARδ. Both LA and PPARδ have been previously and independently shown to play a role in normal and cancer SC self-renewal, both in hematopoietic stem cells and epithelial tumors like colorectal cancer ^29,30^. We expand these results by showing that in HSPC LA activity is specifically exerted on PPARδ, by enhancing its binding to chromatin and activating a specific transcriptional programme aimed at regulating lipid metabolism, and that the functional consequences of this interaction are enhanced by PML-RARAα.

The specific molecular mechanism by which PML-RARα enhances LA activity on PPARδ remains to be fully understood. Both wild type PML and RARα are known to functionally interact with PPARδ, since PML is required for adequate PPARδ signaling and HSC self-renewal ^30^, and PPARs and RARα share RXR as a transcriptional co-factor ^34^. Functional interaction between PML-RARα and PPARδ may occur at several levels. At the level of proximal signaling, the rate of conversion of LA into PPARδ agonists may be enhanced in APLs due to PML-RARα direct or indirect upregulation of lipoxygenase and other LA-metabolizing genes. At the level of transcriptional regulation, PML-RARα may redistribute the RXR pool making it more accessible to further interaction with PPARδ and possibly other transcription factors involved in fatty acid metabolism. Furthermore, we cannot rule out a possible involvement of PPARα, which, at variance with PPARg, is also activated by LA ^28^. PPARα and PPARδ share high homology in their functional domains (DNA and ligand-binding) and their specificity of action is mainly dictated by their pattern of expression, which has been mostly studied in normal, nontumoral tissues ^35^. PPARα is highly expressed in tissues that are central for systemic-metabolism control, like liver and skeletal muscle, whereas PPARδ has a more ubiquitous distribution.

One key issue to be addressed in future research is whether the modifications imparted by HFD and LA are reversible upon subsequent dietary changes. Our results are not conclusive in this respect, since hematopoietic progenitors from HFD-fed pre-leukemic mice maintained their phenotype (increased self-renewal) during *in vitro* culture, suggesting that the effects of HFD remain upon removal of the HFD metabolic-environment, while the effect of LA on self-renewal of progenitors *in vitro* required continuous LA stimulation to be preserved. Possibly, the chronic stimulation achieved through long-term HFD, as opposed to the acute stimulation achieved by short-term in vitro LA exposure, may induce epigenetically stable changes in the transcriptome that render HPCs hypersensitive to fatty acids contained in *in vitro* media. Ongoing efforts are aimed at evaluating whether restrictive dietary-modifications are able to impact on APL development and subsequent growth.

Perhaps surprisingly, we failed to observe clear cooperating genetic events induced by PML-RAR expression and HFD in virtually all APL exomes analysed and in preleukemic cells. Although several studies have shown that additional genetic events in key driver genes lead to accelerated leukemia development in the mouse and poorer outcome in human APLs, the finding that APLs in mice and humans ^5,36^ often lack additional mutations in *bona fide* drivers argue against the invariant need of additional genetic events for APL development. HFD is indeed associated with increased oxidative DNA damage in HSCs, resulting in a moderate increase in mutation rate, yet this does not translate into a higher rate of cooperating mutations, as the few that we found in the HFD established-leukemias and preleukemic cells were not enriched in genes likely to play a causative role. Although our model is not designed to inform on the impact of HFD on the foundational event in APL, usually a translocation involving RARα, we observed no significant effects of HFD on the rate of de novo large scale aberrations in PML-RARα -expressing pre-leukemic stem-cell/progenitors, suggesting that translocations are not per se favoured by HFD-associated oxidative damage. It must be noted, however, that our data do not rule out the possibility that obesity may instead favor the selection of mutations with more aggressive phenotypes at later stages, once the leukemia is fully developed, which may explain its impact on disease outcome rather than risk ^4^. Indeed, our previous analyses of several APL cohorts revealed an enrichment in obese subjects for FLT3 mutations ^3^ (well known to preferentially occur as a late, subclonal event ^37^), a phenomenon that is currently under mechanistic investigation.

Our study also shows that the low-grade, chronic metabolic stress experienced during HFD induces DNA damage-associated exit from quiescence, so far demonstrated only for acute stressors like irradiation, infections and repeated bleeding ^16,38^. Whether DNA damage is a cause or a consequence of the quiescence exit remains a matter of debate. Our data, however, provide a formal demonstration that, at least during HFD, oxidative DNA per se is not sufficient to trigger exit from stem cell quiescence and is likely its consequence, since NAC administration abolished HFD-induced DNA damage but did not affect HFD-induced expansion of the HPC pool. Our work has consequences for informing dietary or pharmacological interventions aimed at counteracting the cancer-promoting effect of obesity. Most importantly, the predominance of nongenetic effects and the reversibility of LA action (at least *in vitro*) suggest that the leukemogenic activity of obesity is partially reversible. Thus, individuals with a history of chronic obesity may still significantly reduce their risk by switching to a healthier lifestyle. This conclusion is supported in other cancers by several retrospective nonrandomized studies ^39,40^, while data on hematological malignances are still missing. Furthermore, we identify LA and probably PUFAs in general as a major target for intervention. LA is the most highly consumed fatty acid in Western diets ^41^; maintaining an adequate ratio of mono-vs poly-unsaturated fatty acids is already a mainstay of several nutritional guidelines, mostly based on epidemiological evidence correlating a high poly- vs mono-unsaturated fats with poor cardiovascular risk. Our data extend the possible utility of these recommendations to leukemia prevention and strengthen their biological rationale. Whether these findings can be generalised to other types of cancer that are heavily influenced by obesity remains to be demonstrated, but we would argue that adequate modeling in the mouse can provide crucial information to increase our understanding of molecular mechanisms, whether common or private to specific cancer types, helping to identify dietary or pharmacological interventions to improve control of what is increasingly perceived as a public health priority.

## Materials and methods

### Data availability

All sequencing data are available as a NCBI Bioproject, n PRJNA812866

### *In vivo* response to HFD

PML-RARA α knock-in mice ^11^ were provided by T.J. Ley and backcrossed in C57BL/6J background for at least 10 generations. Animal received ad libitum food and drinking water and were kept in a regimen of 12-hour light-and-darkness cycle. Animals were fed with ad libitum SD or HFD (Broogarden, D12492) starting from 8-12 weeks of age.

For Comet assays, transplantation, colony formation, sequencing and Immunofluorescence experiments, see supplementary methods

## Acknowledgments

Research in LM lab is funded by AIRC-Cariplo Foundation (TRIDEO 2014 n. 15650), Italian Ministry of Health - Giovani Ricercatori (GR-2011-02350249), European Hematology Association-Jose Carreras Young Investigator Award 2014.

Research in FB lab is funded by AIRC IG20109 and Italian Ministry of Health grants

Research in PGP’s lab is funded by AIRC IG 2017 n. 20162 grant

EGD is partially funded by Ayudas a la Recualificación del sistema universitario español 2021, María Zambrano, Ministerio de Universidades, Spain

We wish to thank colleagues at the former Mario Negri Sud institute for hosting some of the *in vivo* experiments

## Authorship contribution

LM and PGP conceived the study; LM and PF designed and performed *in vivo* and *in vitro* experiments; LM, MA, GT, EG, EB, MB, SR, CR performed bioinformatic analyses; RP and GID designed and performed ChIPseq; EGD, DV, AG, BG, SO, BAD performed *in vitro* experiments; TD, RP, AS participated to *in vivo* experiments; MS, FB, MA, GID participated in the design of the experiments; LM, PF and PGP wrote the manuscript; all authors revised the manuscript

## Supplementary figures and tables

**Supplementary figure 1**

A: mutations by age divided in 3 BMI classes

B: copy number alterations as analysed by CNVkit

**Supplementary figure 2**

A. Expression of PPARs in NB4 (RPKM)

B. Distribution of genomic features in PPARδ peaks

C. Odds Ratio vs -log(pVal) of enriched KEGG pathways in PPARδ-associated genes

Supplementary table 1: BRASS results

Supplementary table 2: KDM6A_copies

Supplementary table 3: APL and colonies genes, VEP-annotated

Supplementary table 4: Differentially expressed genes between LINO- and BSA-treated NB4 cells

Supplementary table 5: KEGG GSEA LINO- vs BSA-treated NB4 cells

Supplementary table 6: KEGG enrichR LINO- vs BSA-treated NB4 cells

Supplementary table 7: PPARd peaks

Supplementary table 8: PPARD enricheR KEGG

Supplementary table 9: PPARD enriched motifs

## References

1. Avgerinos KI, Spyrou N, Mantzoros CS, Dalamaga M. Obesity and cancer risk: Emerging biological mechanisms and perspectives. Metabolism. 2019;92:121–135.

2. Bhaskaran K, Douglas I, Forbes H, et al. Body-mass index and risk of 22 specific cancers: a population-based cohort study of 5·24 million UK adults. Lancet. 2014;384(9945):755–765.

3. Mazzarella L, Botteri E, Matthews A, et al. Obesity is a risk factor for acute promyelocytic leukemia: evidence from population and cross-sectional studies studies and correlation with flt3 mutations and polyunsaturated fatty acid metabolism. Haematologica. 2019;

4. Breccia M, Mazzarella L, Bagnardi V, et al. Increased BMI correlates with higher risk of disease relapse and differentiation syndrome in patients with acute promyelocytic leukemia treated with the AIDA protocols Increased BMI correlates with higher risk of disease relapse and differentiation syndro. 2013;119(1):49–54.

5. Ronchini C, Brozzi A, Riva L, et al. PML-RARA-associated cooperating mutations belong to a transcriptional network that is deregulated in myeloid leukemias. Leukemia. 2017;31(9):1975–1986.

6. Cole CB, Verdoni AM, Ketkar S, et al. PML-RARA requires DNA methyltransferase 3A to initiate acute promyelocytic leukemia. J. Clin. Invest. 2016;126(1):85–98.

7. Wartman LD, Larson DE, Xiang Z, et al. Sequencing a mouse acute promyelocytic leukemia genome reveals genetic events relevant for disease progression. J. Clin. Invest. 2011;121(4):1445–1455.

8. Leukemia AM. Genomic and epigenomic landscapes of adult de novo acute myeloid leukemia. N. Engl. J. Med. 2013;368(22):2059–74.

9. Nasr R, Guillemin M-C, Ferhi O, et al. Eradication of acute promyelocytic leukemia-initiating cells through PML-RARA degradation. Nat. Med. 2008;14(12):1333–1342.

10. Kelly LM, Kutok JL, Williams IR, et al. PML/RARalpha and FLT3-ITD induce an APL-like disease in a mouse model. Proc Natl Acad Sci U S A. 2002;99(12):8283–8288.

11. Westervelt P, Lane AA, Pollock JL, et al. High-penetrance mouse model of acute promyelocytic leukemia with very low levels of PML-RARalpha expression. Blood. 2003;102(5):1857–1865.

12. Hariri N, Thibault L. High-fat diet-induced obesity in animal models. Nutr. Res. Rev. 2010;23(2):270–299.

13. Bankoglu EE, Seyfried F, Arnold C, et al. Reduction of DNA damage in peripheral lymphocytes of obese patients after bariatric surgery-mediated weight loss. Mutagenesis. 2018;33(1):61–67.

14. Kompella P, Vasquez KM. Obesity and cancer: A mechanistic overview of metabolic changes in obesity that impact genetic instability. Mol. Carcinog. 2019;58(9):1531–1550.

15. Beerman I, Seita J, Inlay MA, Weissman IL, Rossi DJ. Quiescent hematopoietic stem cells accumulate DNA damage during aging that is repaired upon entry into cell cycle. Cell Stem Cell. 2014;15(1):37–50.

16. Insinga A, Cicalese A, Faretta M, et al. DNA damage in stem cells activates p21, inhibits p53, and induces symmetric self-renewing divisions. Proc. Natl. Acad. Sci. U. S. A. 2013;110(10):3931–6.

17. Wartman L, Larson D, Xiang Z. Sequencing a mouse acute promyelocytic leukemia genome reveals genetic events relevant for disease progression. J. Clin. 2011;121(4):1445–1455.

18. McLaren W, Gil L, Hunt SE, et al. The Ensembl Variant Effect Predictor. Genome Biol. 2016;17(1):122.

19. Bailey MH, Tokheim C, Porta-Pardo E, et al. Comprehensive Characterization of Cancer Driver Genes and Mutations. Cell. 2018;173(2):371–385.e18.

20. Azmi AS, Uddin MH, Mohammad RM. The nuclear export protein XPO1 — from biology to targeted therapy. Nat. Rev. Clin. Oncol. 2021;18(3):152–169.

21. Whale AD, Colman L, Lensun L, Rogers HL, Shuttleworth SJ. Functional characterization of a novel somatic oncogenic mutation of PIK3CB. Signal Transduct. Target. Ther. 2017;2(1):17063.

22. Degos L, Wang ZY. All trans retinoic acid in acute promyelocytic leukemia. Oncogene. 2001;20(49):7140–7145.

23. Wojiski S, Guibal FC, Kindler T, et al. PML–RARα initiates leukemia by conferring properties of self-renewal to committed promyelocytic progenitors. Leukemia. 2009;23(8):1462–1471.

24. Lee J-M, Govindarajah V, Goddard B, et al. Obesity alters the long-term fitness of the hematopoietic stem cell compartment through modulation of Gfi1 expression. J. Exp. Med. 2018;215(2):627–644.

25. Oguro H, Ding L, Morrison SJ. SLAM family markers resolve functionally distinct subpopulations of hematopoietic stem cells and multipotent progenitors. Cell Stem Cell. 2013;13(1):102–116.

26. Samudio I, Harmancey R, Fiegl M, et al. Pharmacologic inhibition of fatty acid oxidation sensitizes human leukemia cells to apoptosis induction. J. Clin. Invest. 2010;120(1):142–156.

27. Forman BM, Chen J, Evans RM. Hypolipidemic drugs, polyunsaturated fatty acids, and eicosanoids are ligands for peroxisome proliferator-activated receptors alpha and delta. Proc. Natl. Acad. Sci. U. S. A. 1997;94(9):4312–4317.

28. Kliewer SA, Sundseth SS, Jones SA, et al. Fatty acids and eicosanoids regulate gene expression through direct interactions with peroxisome proliferator-activated receptors α and γ. Proc. Natl. Acad. Sci. 1997;94(9):4318 LP–4323.

29. Beyaz S, Mana MD, Roper J, et al. High-fat diet enhances stemness and tumorigenicity of intestinal progenitors. Nature. 2016;531(7592):53–58.

30. Ito K, Carracedo A, Weiss D, et al. A PML–PPAR-δ pathway for fatty acid oxidation regulates hematopoietic stem cell maintenance. Nat. Med. 2012;18(9):1350–8.

31. Lalevée S, Anno YN, Chatagnon A, et al. Genome-wide in silico identification of new conserved and functional retinoic acid receptor response elements (direct repeats separated by 5 bp). J. Biol. Chem. 2011;286(38):33322–33334.

32. Shearer BG, Wiethe RW, Ashe A, et al. Identification and characterization of 4-chloro-N-(2-{[5-trifluoromethyl)-2-pyridyl]sulfonyl}ethyl)benzamide (GSK3787), a selective and irreversible peroxisome proliferator-activated receptor delta (PPARdelta) antagonist. J. Med. Chem. 2010;53(4):1857–1861.

33. Welch JS, Yuan W, Ley TJ. PML-RARA can increase hematopoietic self-renewal without causing a myeloproliferative disease in mice. J. Clin. Invest. 2011;121(4):1636–1645.

34. Kliewer SA, Umesono K, Noonan DJ, Heyman RA, Evans RM. Convergence of 9-cis retinoic acid and peroxisome proliferator signalling pathways through heterodimer formation of their receptors. Nature. 1992;358(6389):771–774.

35. Berger J, Moller DE. The Mechanisms of Action of PPARs. Annu. Rev. Med. 2002;53(1):409–435.

36. Lehmann-Che J, Bally C, Letouzé E, et al. Dual origin of relapses in retinoic-acid resistant acute promyelocytic leukemia. Nat. Commun. 2018;9(1):2047.

37. Papaemmanuil E, Gerstung M, Bullinger L, et al. Genomic Classification and Prognosis in Acute Myeloid Leukemia. N. Engl. J. Med. 2016;374(23):2209–2221.

38. Walter D, Lier A, Geiselhart A, et al. Exit from dormancy provokes DNA-damage-induced attrition in haematopoietic stem cells. Nature. 2015;520(7548):549–552.

39. Teras LR, Patel A V, Wang M, et al. Sustained Weight Loss and Risk of Breast Cancer in Women 50 Years and Older: A Pooled Analysis of Prospective Data. J. Natl. Cancer Inst. 2020;112(9):929–937.

40. Ligibel JA, Alfano CM, Courneya KS, et al. American Society of Clinical Oncology position statement on obesity and cancer. J. Clin. Oncol. Off. J. Am. Soc. Clin. Oncol. 2014;32(31):3568–3574.

41. Rett BS, Whelan J. Increasing dietary linoleic acid does not increase tissue arachidonic acid content in adults consuming Western-type diets: a systematic review. Nutr. Metab. (Lond). 2011;8(1):36.

42. Grignani F, Kinsella T, Mencarelli A, et al. High-efficiency gene transfer and selection of human hematopoietic progenitor cells with a hybrid EBV/retroviral vector expressing the green fluorescence protein. Cancer Res. 1998;58(1):14–19.

43. Kim D, Pertea G, Trapnell C, et al. TopHat2: accurate alignment of transcriptomes in the presence of insertions, deletions and gene fusions. Genome Biol. 2013;14(4):R36.

44. Anders S, Pyl PT, Huber W. HTSeq-A Python framework to work with high-throughput sequencing data. Bioinformatics. 2015;31(2):166–169.

45. Robinson MD, McCarthy DJ, Smyth GK. edgeR: a Bioconductor package for differential expression analysis of digital gene expression data. Bioinformatics. 2010;26(1):139–140.

46. Li H, Durbin R. Fast and accurate long-read alignment with Burrows-Wheeler transform. Bioinformatics. 2010;26(5):589–95.

47. Cibulskis K, Lawrence MS, Carter SL, et al. Sensitive detection of somatic point mutations in impure and heterogeneous cancer samples. Nat Biotech. 2013;31(3):213–219.

48. Nik-Zainal S, Davies H, Staaf J, et al. Landscape of somatic mutations in 560 breast cancer whole-genome sequences. Nature. 2016;534(7605):47–54.

49. McKenna A, Hanna M, Banks E, et al. The Genome Analysis Toolkit: a MapReduce framework for analyzing next-generation DNA sequencing data. Genome Res. 2010;20(9):1297–303.

50. Talevich E, Shain AH, Botton T, Bastian BC. CNVkit: Genome-Wide Copy Number Detection and Visualization from Targeted DNA Sequencing. PLoS Comput. Biol. 2016;12(4):e1004873.

51. Conway JR, Lex A, Gehlenborg N. UpSetR: an R package for the visualization of intersecting sets and their properties. Bioinformatics. 2017;33(18):2938–2940.

52. Dellino GI, Palluzzi F, Chiariello AM, et al. Release of paused RNA polymerase II at specific loci favors DNA double-strand-break formation and promotes cancer translocations. Nat. Genet. 2019;51(6):1011–1023.

53. Li H, Handsaker B, Wysoker A, et al. The Sequence Alignment/Map format and SAMtools. Bioinforma. Appl. NOTE. 2009;25(16):2078–2079.

54. Zhang Y, Liu T, Meyer C, Eeckhoute J, Johnson D. Model-based analysis of ChIP-Seq (MACS). Genome Biol. 2008;9:.

55. Heinz S, Benner C, Spann N, et al. Simple combinations of lineage-determining transcription factors prime cis-regulatory elements required for macrophage and B cell identities. Mol. Cell. 2010;38(4):576–589.

56. Ramírez F, Ryan DP, Grüning B, et al. deepTools2: a next generation web server for deep-sequencing data analysis. Nucleic Acids Res. 2016;44(W1):W160–W165.

