## Supplementary figures and images for "High-fat diet promotes Acute Promyelocytic Leukemia through PPARδ-enhanced self-renewal of preleukemic progenitors"

### supplementary figure 1

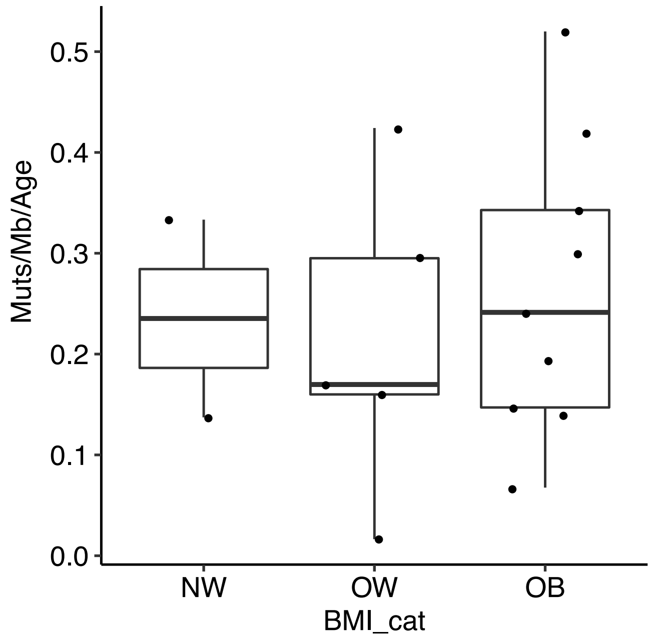

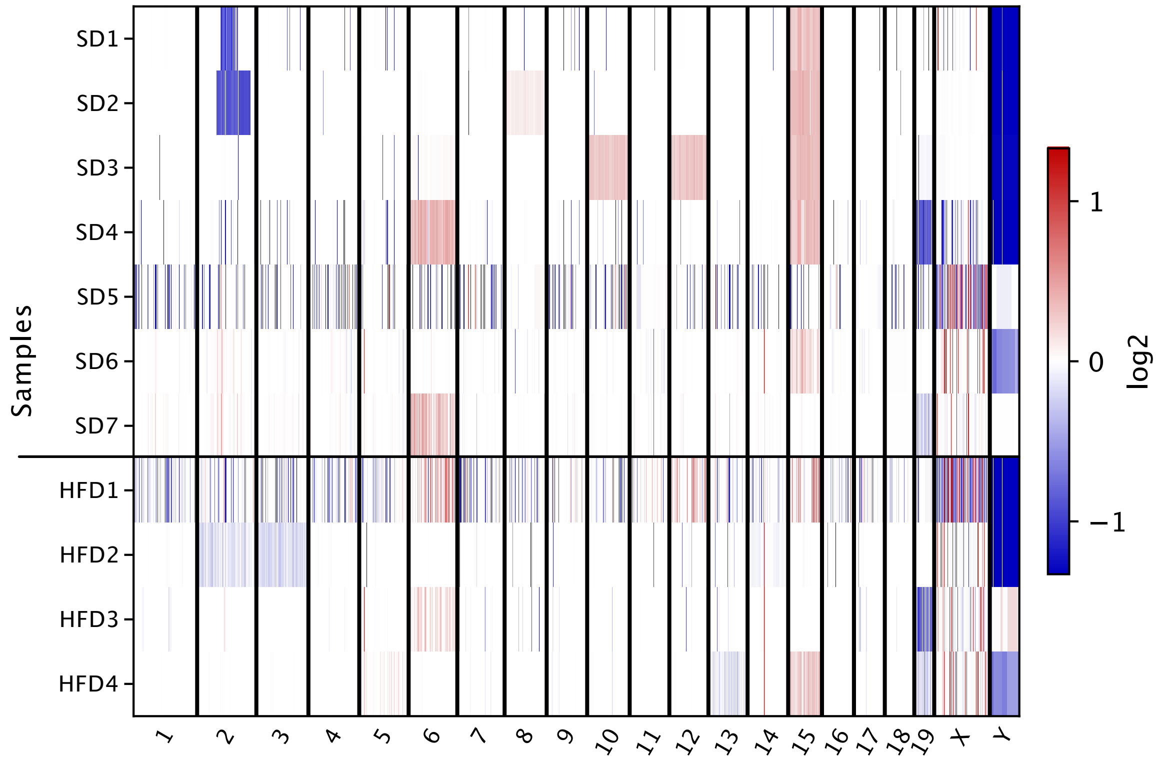

### supplementary figure 2

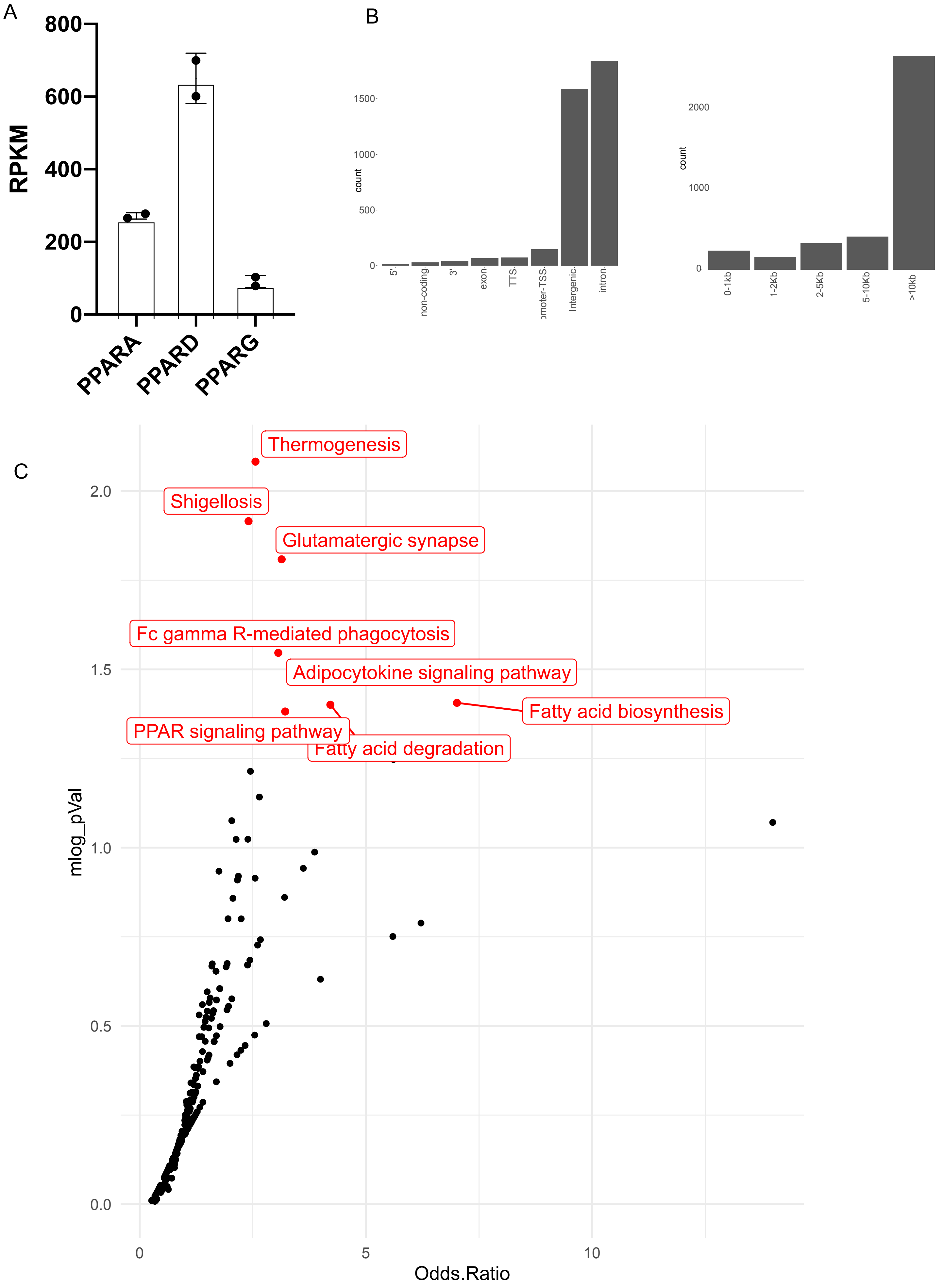
